# FTO suppresses STAT3 activation and modulates proinflammatory interferon-stimulated gene expression

**DOI:** 10.1101/2021.07.23.453596

**Authors:** Michael J. McFadden, Matthew T. Sacco, Kristen A. Murphy, Moonhee Park, Nandan S. Gokhale, Kim Y. Somfleth, Stacy M. Horner

## Abstract

Signaling initiated by type I interferon (IFN) results in the induction of hundreds of IFN-stimulated genes (ISGs). The type I IFN response is important for antiviral restriction, but aberrant activation of this response can lead to inflammation and autoimmunity. Regulation of this response is incompletely understood. We previously reported that the mRNA modification m^6^A and its deposition enzymes, METTL3 and METTL14 (METTL3/14), promote the type I IFN response by directly modifying the mRNA of a subset of ISGs to enhance their translation. Here, we determined the role of the RNA demethylase FTO in the type I IFN response. FTO, which can remove either m^6^A or the cap-adjacent m^6^Am RNA modifications, has previously been associated with obesity and body mass index, type 2 diabetes, cardiovascular disease, and inflammation. We found that FTO suppresses the transcription of a distinct set of ISGs, including many known pro-inflammatory genes, and that this regulation is not through the actions of FTO on m^6^Am. Further, we found that depletion of FTO led to activation of STAT3, a transcription factor that mediates responses to various cytokines, but whose role in the type I IFN response is not well understood. This activation of STAT3 increased the expression of a subset of ISGs. Importantly, this increased ISG induction resulting from FTO depletion was partially ablated by depletion of STAT3. Together, these results reveal that FTO negatively regulates STAT3-mediated signaling that induces proinflammatory ISGs during the IFN response, highlighting an important role for FTO in suppression of inflammatory genes.

## Introduction

The type I interferon (IFN) response induces the expression of hundreds of IFN-stimulated genes (ISGs), many of which have antiviral and proinflammatory functions, and thus is crucial for the early response to viral infection [1]. Type I IFNs signal through a dimeric receptor composed of IFNAR1 and IFNAR2 to activate the transcription factors STAT1 and STAT2, which heterodimerize with IRF9 to form the transcription factor complex ISGF3 [2]. ISGF3 then binds to the promoters of ISGs to induce their transcription, resulting in the establishment an antiviral cellular state [3]. While transcriptional induction of ISGs by ISGF3 is the primary driver of the type I IFN response, regulation of this response by other transcription factors or by post-transcriptional controls is incompletely understood. Specifically, the functions of the transcription factor STAT3 in the type I IFN response are not clear, although it has been shown to have functions either in suppressing this response or promoting the expression of certain ISGs, likely in a cell type-specific fashion [4]. We previously established that the addition of the mRNA modification m^6^A to a subset of ISGs by the m^6^A-methyltransferase complex proteins METTL3 and METTL14 (METTL3/14) increases their translation by recruiting the m^6^A-binding protein YTHDF1 and that this promotes an antiviral cellular state [5]. However, the role of the m^6^A demethylase FTO in the type I IFN response has not yet been described. The functions of FTO are important for several aspects of human health, and genetic variants of this gene are associated with obesity and body mass index, type 2 diabetes, cardiovascular disease, and inflammation [6–9]. Despite its essential functions for human development and health [10], the mechanisms by which FTO regulates these biological processes are incompletely understood.

A major molecular function of FTO, which has sequence homology to alpha-ketoglutarate oxygenase enzymes such as the AlkB homolog family (ALKBH) proteins, is RNA demethylation [11]. Importantly, FTO has been shown to be involved in the demethylation of both m^6^A [12], a function shared by ALKBH5 [13], and the cap-adjacent m^6^Am modification [14]. m^6^A is deposited by a complex of proteins, including METTL3/14 and others [15], and can regulate many aspects of RNA metabolism, including degradation and translation, among other processes [16–19]. m^6^Am addition is catalyzed by the enzyme PCIF1 at the first transcribed nucleotide in an mRNA and may be involved in regulating mRNA stability and translation [20–22]. Through its RNA demethylase functions, FTO can regulate many cellular and biological processes, including neurogenesis, dopamine signaling and appetite regulation, adipogenesis, oncogenesis, and viral infection [23–29]. Interestingly, FTO depletion was recently shown to sensitize melanoma cells to IFN-γ treatment, suggesting a role for FTO in the response to IFNs [28]. Further, FTO has been shown to regulate infection by a number of viruses, presumably by demethylating viral RNA [29], but its role in regulation of host responses to viral infection has not been elucidated. Therefore, we sought to define the role of FTO in the type I IFN response.

Here, we found that depletion of FTO results in increased production of a subset of ISGs. However, whereas METTL3/14 promotes the translation of a subset of ISGs without affecting their mRNA levels [5], FTO regulates a distinct subset of ISGs at the mRNA level. By labeling nascent RNA, we found that FTO inhibits the transcription of these ISGs and that FTO-depleted cells are primed to respond to type I IFN. FTO regulation of ISGs is not through demethylation of m^6^Am, as deletion of the m^6^Am writer enzyme PCIF1 did not impact the phenotypic effect of FTO depletion on ISG expression. Interestingly, we found that FTO depletion results in increased phosphorylation and thus activation of transcription factor STAT3. Additionally, STAT3 activation through treatment with the cytokine IL6 or expression of the *Salmonella* effector protein SarA, known to induce STAT3 phosphorylation, recapitulated the phenotypic effects of STAT3 activation by FTO depletion [30, 31], suggesting that suppression of STAT3 activation by FTO represses transcription of a specific subset of ISGs. In support of this hypothesis, depletion of STAT3 led to partial ablation of FTO-mediated suppression of ISG induction. Taken together, these results reveal a novel role for FTO in the transcriptional suppression of STAT3-regulated ISGs, which has important implications for our understanding of the role of FTO in disease.

## Results

### FTO regulates the mRNA expression of a subset of ISGs

Having previously shown that the m^6^A-methyltransferase complex proteins METTL3/14 promote the translation of specific ISGs via the addition of m^6^A to these ISGs [5], we hypothesized that the major m^6^A-demethylase FTO would suppress the translation of these ISGs by removal of m^6^A. To test this hypothesis, we used siRNAs to deplete METTL3/14 or FTO in Huh7 cells and induced the expression of ISGs by treatment with IFN-β, a type I IFN. Similar to our previous results, depletion of METTL3/14 resulted in reduced protein levels of IFITM1, GBP1, IFIT3, and MX1 [5]. Depletion of FTO showed the opposite result in that its depletion resulted in increased protein expression of the METTL3/14-regulated ISGs, expect for MX1 whose expression was only modestly affected by METTL3/14 depletion and not altered by FTO depletion (Figure 1A). As before, the ISGs ISG15 and EIF2AK2 were unaffected by METTL3/14 depletion, and here we also found that their IFN-induced expression was unaffected by FTO depletion (Figure 1A). We next determined whether FTO depletion changed the mRNA induction of these ISGs. Interestingly, while METTL3/14 depletion did not impact the mRNA levels of these ISGs, as expected [5], FTO depletion did increase the mRNA levels of the regulated ISGs, including *IFITM1*, *GBP1*, and *IFIT3* (Figure 1B). These results show that FTO negatively regulates specific ISGs at the mRNA level. Thus, the mechanism underlying the regulatory role of FTO on ISGs is distinct from METTL3/14.

**Figure 1:**
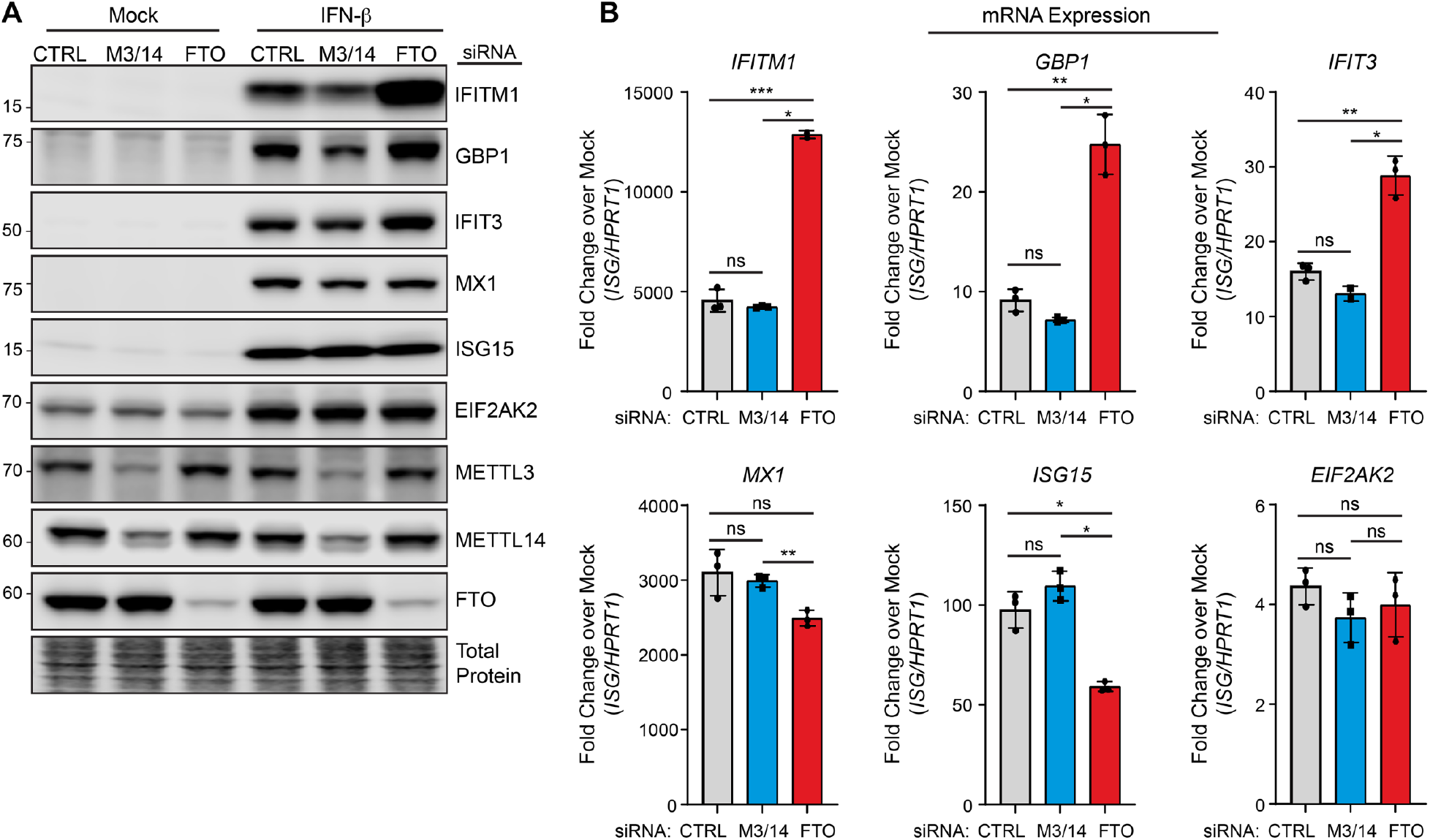
FTO regulates the mRNA expression of a subset of ISGs. **(A)** Immunoblot analysis of extracts from Huh7 cells transfected with siRNAs to control (CTRL), METTL3/14 (M3/14), or FTO prior to mock or IFN-β (12 h) treatment. Data are representative of 3 biological experiments. **(B)** RT-qPCR analysis of ISG induction normalized to *HPRT1* following IFN-β treatment (12 hours) of Huh7 cells treated with non-targeting control (CTRL) or METTL3/14 siRNA plotted as fold change over mock treatment for each ISG. Values are the mean ± SD of 3 technical replicates, representative of 3 biological experiments. * p < 0.05, ** p < 0.01, *** p < 0.005 by one way ANOVA with Dunnett’s multiple comparisons test.

### FTO regulates the transcription of certain ISGs

To determine how FTO regulates the mRNA expression of ISGs, we pulsed siCTRL, siMETTL3/14, or siFTO-treated cells with 4-thiouridine (4-sU) and IFN-β for 1 hour to metabolically label nascent transcripts produced by IFN stimulation. We then purified the nascent RNA and performed RT-qPCR to quantify relative transcription. These experiments revealed that depletion of FTO led to a marked increase of the transcription of the FTO-regulated ISG *IFITM1* during the pulse. Interestingly, the transcription of the non-FTO-regulated ISGs *ISG15* and *EIF2AK2* were also slightly increased by FTO depletion (Figure 2A). METTL3/14 depletion did not affect the transcription of these ISGs nor a known m^6^A-modified gene, *CREBBP*, whose stability is regulated by m^6^A [16] (Figure 2A). Next, to determine how FTO regulates the transcription of ISGs over time, we performed the 4-sU and IFN-β pulses at both 1 and 12 hours in siRNA-treated cells and quantified the transcription of a broader set of ISGs by RT-qPCR (Figure 2B). Similar to the results shown in Figure 2A, following the 1 hour 4-sU and IFN-β pulses nearly all ISGs tested (*IFITM1*, *GBP1*, *IFIT3*, *MX1*, *ISG15*, and *EIF2AK2*) displayed some increased transcription following FTO depletion. However, following the 12 hour 4-sU and IFN-β pulses, only *IFITM1, GBP1, and IFIT3* still had increased transcription, while the non-FTO regulated ISGs *MX1*, *ISG15*, and *EIF2AK2* did not (Figure 1B; Figure 2B). These data indicate that FTO depletion primes cells to respond to IFN-β, as they transcribe ISGs more rapidly, but this early transcriptional upregulation only persist for a subset of ISGs.

**Figure 2:**
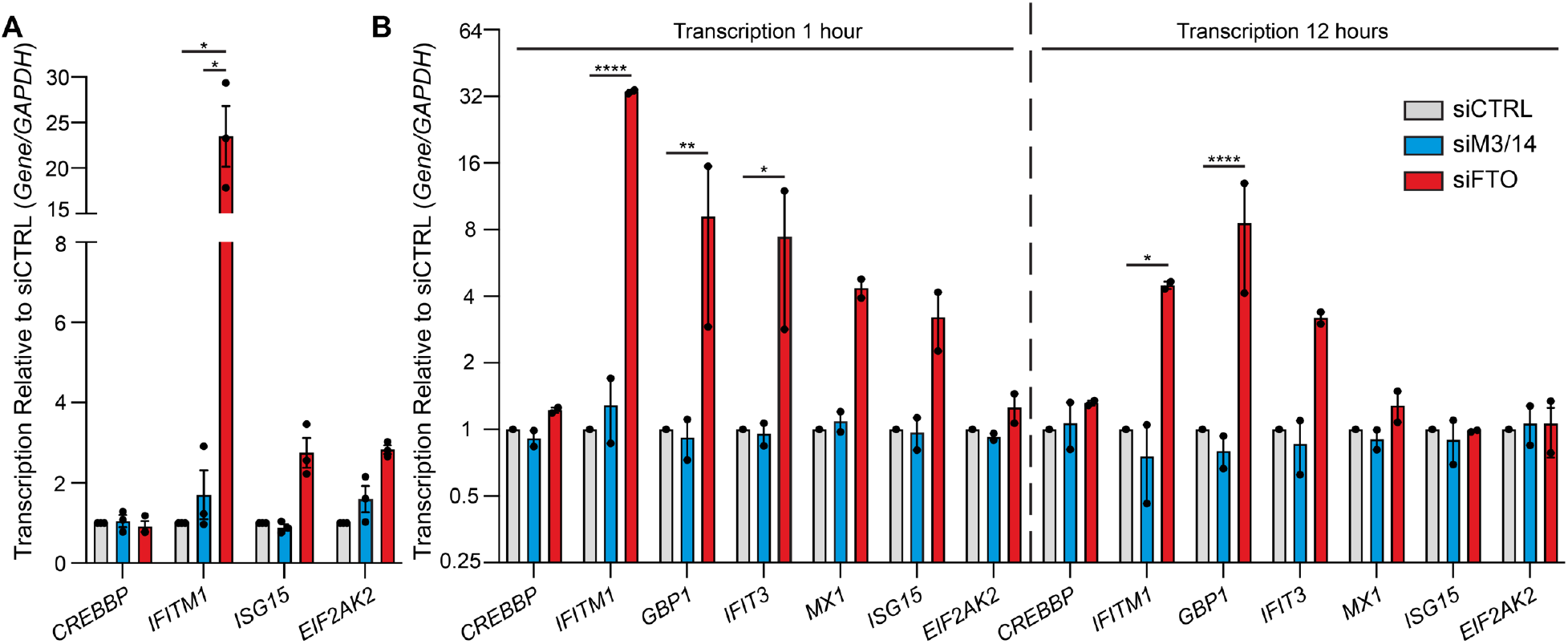
FTO regulates the transcription of ISGs. **(A-B)** Relative transcription analysis of Huh7 cells treated with siRNAs to control (CTRL), METTL3/14 (M3/14), or FTO prior to mock or IFN-β treatment + 4-sU pulse labeling for 1 hour (A) or 1 and 12 hours (B). Values are the mean ± SEM of 3 (A) or 2 (B) biological experiments. * P < 0.01; all other comparisons not significant (P > 0.05) by 2-way ANOVA with Tukey’s multiple comparisons.

### FTO regulation of ISGs occurs independently of m^6^Am

FTO can catalyze the removal of both m^6^A and m^6^Am [12, 14]. To determine if FTO acts on m^6^Am or on m^6^A to suppress the expression of ISGs, we removed expression of the m^6^Am-methylase PCIF1 [20, 21] using PCIF1 knockout cells generated by CRISPR/Cas9 [20]. Then, we tested if loss of m^6^Am by PCIF1 knockout could abrogate suppression of ISGs by FTO. To do this, we depleted FTO in either parental 293T cells or in two independent clonal cell lines in which the m^6^Am-methylase PCIF1 was removed by CRISPR/Cas9 [20]. Following IFN-β treatment for 12 hours, we measured the expression of the FTO-regulated ISGs IFITM1 and IFIT3 by immunoblot and found that the upregulation of IFITM1 and IFIT3 seen upon FTO depletion was not altered by loss of PCIF1 expression (Figure 3A). These results reveal that FTO regulation of ISGs does not occur through m^6^Am modification of RNAs.

**Figure 3:**
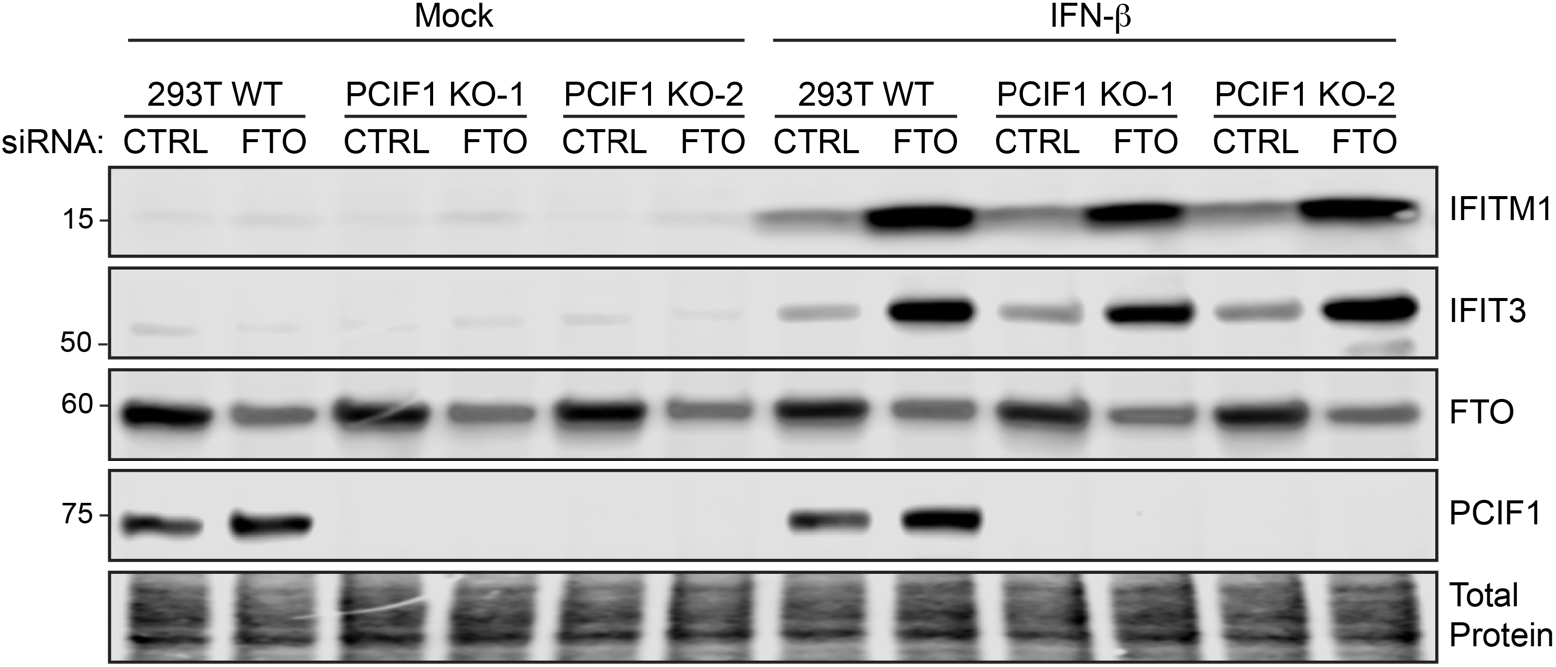
FTO regulation of ISGs occurs independently of m^6^Am. Immunoblot analysis of extracts from wild-type (WT) 293T cells or two clonal PCIF1 knockout cell lines transfected with siRNAs to control (CTRL) or FTO prior to mock or IFN-β (12 h) treatment. Data are representative of 3 biological experiments.

### FTO suppresses inflammatory gene expression

Having determined that FTO regulates the transcription of certain ISGs, we next sought to more broadly characterize the regulatory role of FTO during the type I IFN response. Using RNA-seq, we profiled gene expression in FTO-depleted cells, following mock treatment or IFN-β stimulation (8 hours) (Table S1). Genes regulated by FTO in mock-treated cells were enriched in biological categories such as RNA metabolism and gene expression, supporting the known role of FTO in RNA regulation, as well as immune categories such as response to IFN-γ and response to external biotic stimulus (Table S2.1) [12]. Gene ontology analysis of the genes significantly regulated by FTO (adjusted P < 0.01) during the IFN response revealed that FTO-regulated genes are enriched in biological categories related to immunity, defense responses, and cytokines (Figure 4A; Table S2.2. To determine the effect of FTO depletion on the IFN-induced expression of ISGs, we profiled the 300 most highly-induced ISGs that we had previously defined [5] (Figure 4B; Table S1.1) and found that FTO depletion led to increased expression of a subset of these ISGs (Figure 4C; Table S1.2). Indeed, 140 of these ISGs were significantly regulated by FTO (adjusted P<0.01) and of these 140, nearly all (136) were upregulated following FTO depletion, confirming that FTO is a suppressor of a distinct set of ISGs. While these IGSs are also induced following FTO depletion in the absence of IFN treatment (Figure 4D; Table S1.3), these effects were generally exacerbated by IFN-β treatment, as seen in a heatmap of the top 50 most highly-induced FTO-regulated ISGs (Figure 4E), further supporting the idea that FTO-depleted cells are primed to respond to IFN-β. The ISGs most highly upregulated by FTO depletion include the proinflammatory chemokines *CXCL10* and *CXCL11* (Figure 4A). Indeed, our analysis of proinflammatory gene expression revealed that FTO depletion generally led to an upregulation of proinflammatory gene expression [32], with strong upregulation of the proinflammatory cytokines *TNF* and *IL6* and the proinflammatory chemokines *CXCL9, CXCL10,* and *CXCL11* (Figure 4F). Together, these results reveal that FTO negatively regulates a subset of ISGs, including many proinflammatory genes.

**Figure 4:**
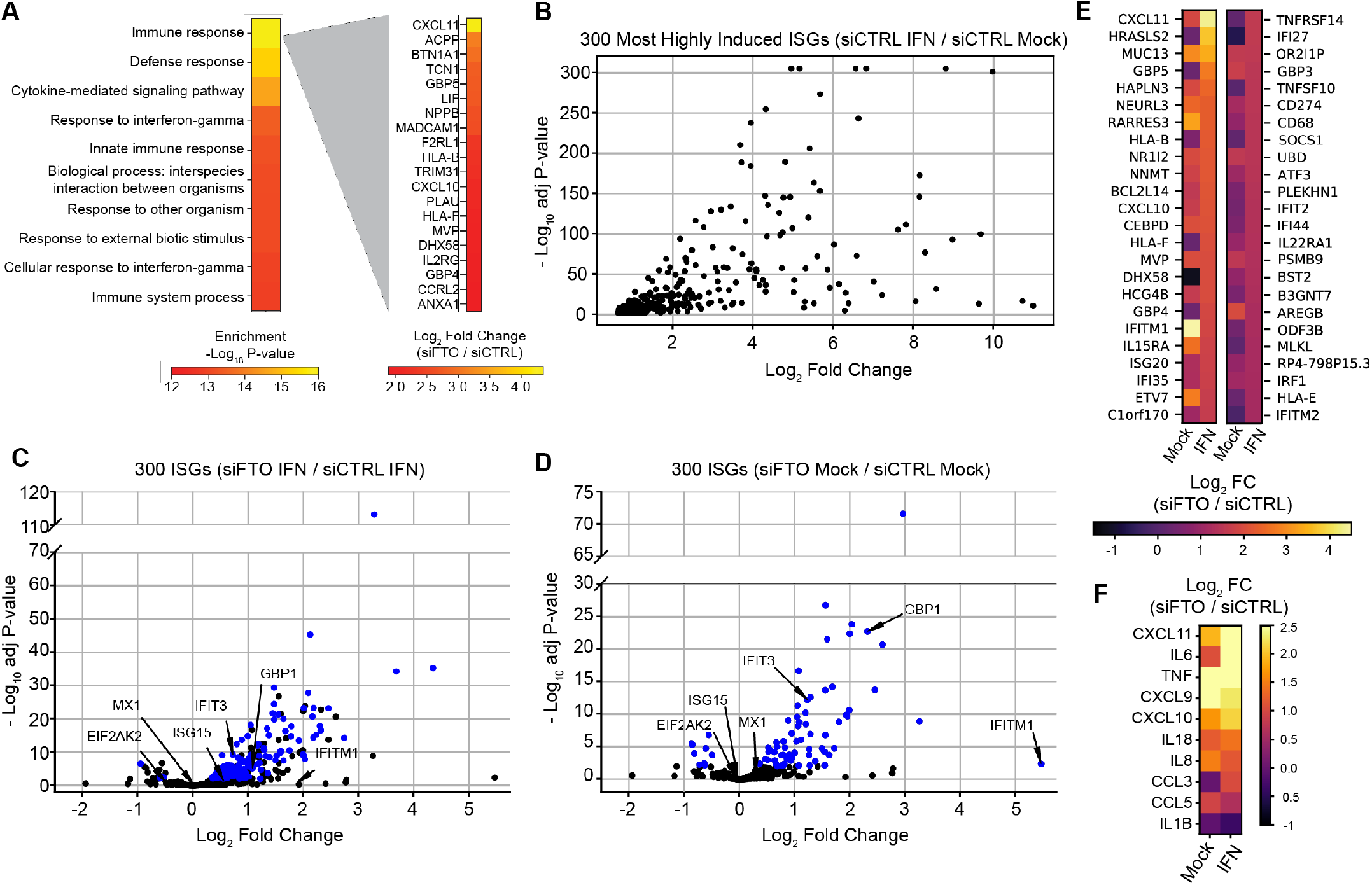
FTO suppresses inflammatory gene expression. RNA-seq analysis of Huh7 cells treated with siRNAs to control (CTRL) or FTO (36 h), followed by mock or IFN-β treatment (8 h). **(A)** Gene ontology analysis of differentially expressed genes in IFN-β-treated cells (siFTO / siCTRL). Left panel shows biological categories and right panel shows differentially expressed genes within the immune response category. **(B-D)** Volcano plots of the 300 most highly-induced ISGs (adjP<0.01; basemean > 50) (A), and the effect of FTO depletion on these 300 ISGs (siFTO / siCTRL) in IFN-β treated cells (B), or mock-treated cells (C). **(E)** Heatmap of the effect of FTO depletion on the 50 most induced ISGs (siFTO / siCTRL) following mock and IFN-β treatment. **(F)** Heatmap of the effect of FTO depletion on canonical proinflammatory cytokines and chemokines (siFTO / siCTRL).

### FTO suppresses STAT3 activation

Having found that FTO suppresses the transcription of only a subset of ISGs and that FTO-depleted cells are primed to respond to type I IFN, we hypothesized that FTO may regulate these ISGs through a transcription factor other than the canonical ISGF3 complex of STAT1, STAT2, and IRF9 [2]. Interestingly, FTO has previously been found to regulate the activation of the transcription factor STAT3 [33], which can drive transcriptional induction of proinflammatory gene expression and possibly certain ISGs [34, 35]. The activation of STAT3 requires its phosphorylation at the tyrosine 705 (Y705) residue [36]. Thus, to test if FTO negatively regulates STAT3 activation we measured phosphorylation of STAT3 at Y705 in METTL3/14- or FTO-depleted Huh7 cells by immunoblot. FTO depletion, but not METTL3/14 depletion, led to an increase in STAT3 phosphorylation, and this increase was similar in the presence or absence of IFN (Figure 5A). To determine if STAT3 activation is sufficient to induce FTO-regulated ISGs, we treated Huh7 cells with IL-6, which induces Y705 phosphorylation of STAT3 [30]. While IL-6 did induce Y705 phosphorylation of STAT3, it did not induce ISG expression, as measured by immunoblot. However, in conjunction with IFN-β treatment, IL-6 did modestly enhance the expression of the FTO-regulated ISGs IFITM1 and IFIT3, but not ISG15 and EIF2AK2. These data suggest in the presence of IFN-β, activation of STAT3 can recapitulate the effect of FTO depletion on ISG expression (Figure 5B). Additionally, STAT3 activation in Huh7 cells by expression of the *Salmonella* effector protein SarA, which is known to activate STAT3 [31], resulted in increased expression of IFITM1 and IFIT3, but not ISG15 and EIF2AK2, following both mock and IFN-β treatment (Figure 5C), supporting our finding that STAT3 activation can regulate ISGs in a similar fashion to FTO depletion during the IFN response. To determine whether STAT3 activation is responsible for the upregulation of ISGs that results from FTO depletion, we co-depleted FTO and STAT3 in Huh7 cells using siRNA. Depletion of STAT3 alone reduced the expression of IFITM1 and IFIT3, but not ISG15 and EIF2AK2 (Figure 5D). Importantly, STAT3 depletion also partially ablated the effect of FTO depletion on IFITM1 and IFIT3 (Figure 5D). Together, these data reveal that FTO suppresses the expression of a subset of ISGs by inhibiting STAT3 activation.

**Figure 5:**
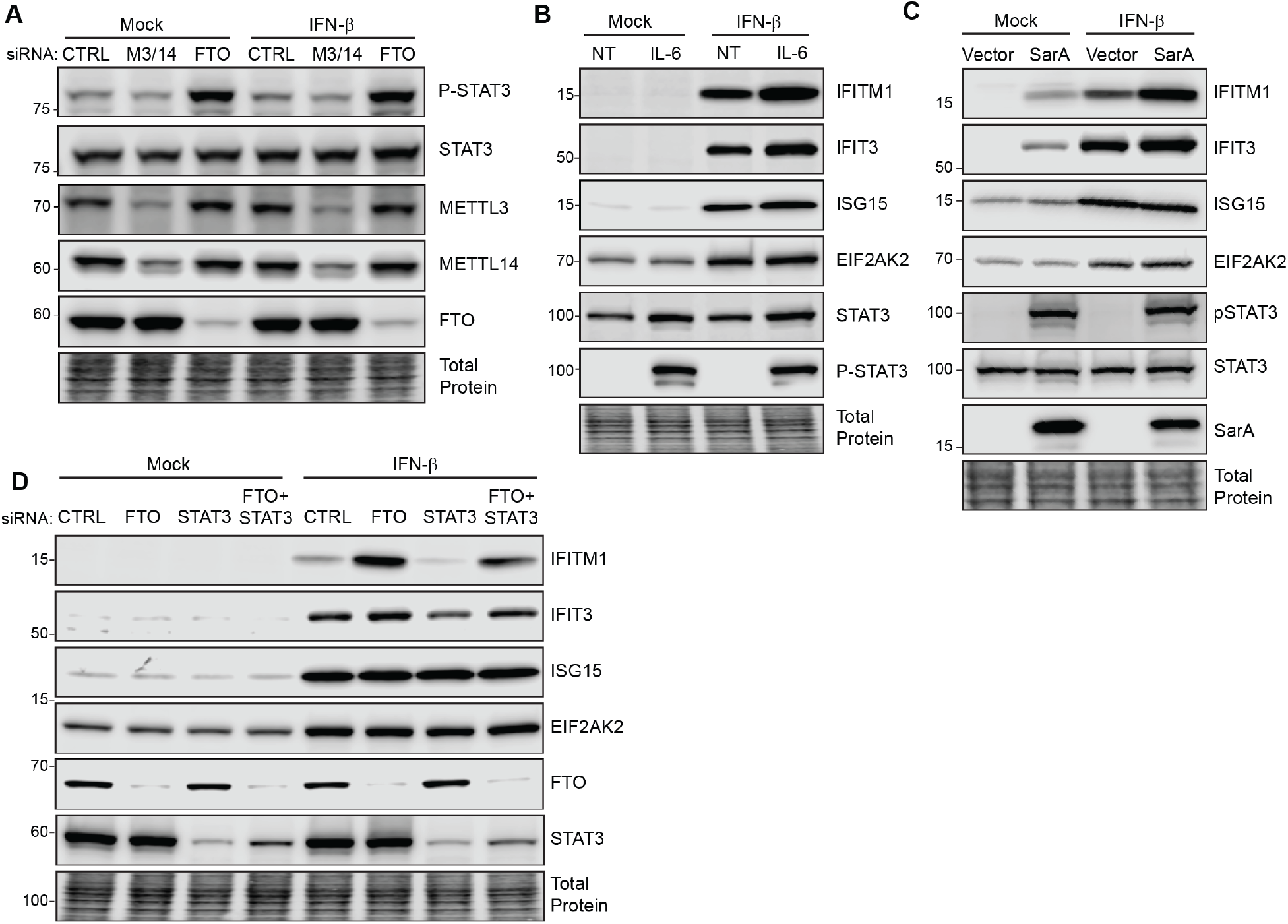
FTO suppresses STAT3 activation. **(A)** Immunoblot analysis of extracts from Huh7 cells transfected with siRNAs to control (CTRL), METTL3/14 (M3/14), or FTO prior to mock or IFN-β (12 h) treatment. **(B)** Immunoblot analysis of extracts from Huh7 treated with mock or IFN-β plus IL-6 or no treatment (NT) for 12 h. **(C)** Immunoblot analysis of extracts from Huh7 cells expressing SarA or vector (16 h), followed by IFN-β (12 h) treatment. **(D)** Immunoblot analysis of extracts from Huh7 cells transfected with siRNAs to control (CTRL), STAT3, FTO, or FTO and STAT3 prior to mock or IFN-β (12 h) treatment. Data in A-D are representative of 3 biological experiments.

## Discussion

We previously found that METTL3/14 and m^6^A facilitate the translation of a subset of m^6^A-modified ISGs [5]. Given the m^6^A and m^6^Am demethylase activities of FTO, we hypothesized that FTO could inhibit the translation of ISGs through demethylation of these modifications on the transcripts of ISGs. However, here, we found that FTO suppresses signaling and transcriptional activation of a subset of ISGs, including many that encode proinflammatory factors. This regulatory function of FTO does not occur through its effect on m^6^Am, as the regulatory effect of FTO on ISGs was intact in cells that do not express the m^6^Am methyltransferase PCIF1. As genetic ablation of METTL3 is not tolerated by Huh7 and other cancer cell lines [37], we were unable to determine whether FTO regulation of ISGs is dependent on METTL3-mediated m^6^A modifications. However, we did find that the major mechanism by which FTO suppressed ISG expression was through its negative regulation of STAT3 activation, which induces specific classes of ISGs (Figure 5). Therefore, our data suggest that active STAT3 may act synergistically with ISGF3 at the promoters of certain ISGs. Given the association of FTO variants with inflammation-related diseases [6–9], these results increase our knowledge of the molecular roles of FTO in human diseases driven by inflammation. Additionally, our work establishes a novel role for FTO as a suppressor of ISGs, and thus defines a new control for the IFN response.

The role of STAT3 in the type I IFN response is unclear. The type I IFN response is typically driven by the ISGF3 transcription factor complex, which is composed of STAT1, STAT2, and IRF9. While this complex does not canonically include STAT3, STAT3 has been shown to be activated by type I IFN in certain cell types [4]. However, most functions of type I IFN signaling, such as antiviral gene expression and immune cell development, are independent of STAT3 [38–40]. In cell types in which STAT3 is activated in response to type I IFN, it has been show to function as either a positive or negative regulator of type I IFN signaling [4, 35, 41]. The mechanisms by which STAT3 may repress type I IFN signaling include sequestration of STAT1 [41], cooperation with repressor proteins such as PLSCR2 [42], or induction of negative regulators of the JAK-STAT pathway, such as SOCS family proteins [43]. However, in addition to these repressive roles, STAT3 has been shown to support or enhance the expression of certain ISGs, such as OAS and MX2 [41], as well as CXCL11 [35], which we found to be among the highest upregulated genes by FTO depletion. Interestingly, in our experiments, STAT3 phosphorylation was not induced by IFN-β treatment alone, but known STAT3 phosphorylation activators such as IL-6 and the Salmonella effector protein SarA induced expression of ISGs that are also increased by FTO depletion (Figure 5). These data suggest that STAT3 can activate the transcription of a subset of ISGs, as well as act synergistically with the ISGF3 transcription factor complex canonically activated by type I IFN to enhance the expression of these ISGs. As such, STAT3 may be responsible for the transcriptional priming of ISGs that we observed following FTO depletion (Figure 2), although this effect seems to be transient for many ISGs, perhaps due to their eventual negative regulation [44]. Whether STAT3 directly promotes ISG transcription by binding to the promoters of these genes or by an indirect mechanism, as well as determining the cell type specificity of this STAT3-mediated regulation, will be an interesting direction for future research.

Previous studies have reported conflicting roles of FTO in regulation of STAT3 activation. In agreement with our findings, FTO overexpression in liver cells can decrease phosphorylation of STAT3 at Y705 and reduce its nuclear localization [33], suggesting that FTO suppresses STAT3 activation. However, other studies in adipocytes have suggested that FTO increases STAT3 activation by stabilizing the mRNA of *JAK2,* which encodes a known STAT3 kinase, to induce STAT3 phosphorylation at Y705 [45]. Therefore, molecular regulation between FTO and STAT3 is complex and may differ in specific cell types. However, given the role of FTO in regulation of mRNA expression, it is likely that FTO regulates the expression of one or more factors that govern STAT3 Y705 phosphorylation through regulation of m^6^A on specific transcripts. Indeed, our RNA-seq data revealed that *JAK2* mRNA expression was upregulated following FTO depletion. As JAK2 encodes a kinase that can phosphorylate STAT3 at Y705 in response to IL6 [46], it is possible that upregulation of JAK2 expression is responsible for the increase in STAT3 phosphorylation and activation observed following FTO depletion. While the m^6^Am demethylase function of FTO is not required for its regulation of ISGs, it is possible that it regulates one or several factors controlling STAT3 activation, such as JAK2, in an m^6^A demethylase-dependent fashion. Indeed, we previously found that JAK2 mRNA is m^6^A-modified during the type I IFN response [5]. However, as FTO depletion may regulate the expression of multiple kinases or phosphatases that control STAT3 phosphorylation [47], additional targeted studies will be required for a more complete understanding of the mechanisms by which FTO regulates STAT3 activation.

In summary, we found that FTO acts as a transcriptional suppressor of a subset of ISGs, including many genes encoding proinflammatory factors. This function of FTO is independent of its role as an m^6^Am demethylase and is mediated at least partially by suppression of STAT3 activation. These results reveal a novel regulatory control over the type I IFN response and shed additional light on the molecular role of FTO in regulation of immune gene expression. As FTO and its substrate modifications m^6^A and m^6^Am can regulate viral infection through their effects on both viral and host RNAs [29, 48, 49], these results will help shape our understanding of the functions of FTO during viral infection. While previous studies have described associations between FTO and immune-related gene expression [27, 50], our work sheds additional light on the mechanisms by which FTO influences innate immune and proinflammatory gene expression. Further, this molecular role of FTO could have important implications for our understanding of the role of FTO in inflammation-related diseases, such as obesity and cancer. Thus, further exploration of the relationship between FTO, STAT3, and inflammatory gene expression in immune cell subsets will be crucial for our understanding of the molecular functions of FTO in disease.

## Acknowledgements

We thank colleagues who provided reagents (see Methods), the Duke Functional Genomics Core Facility, the Duke Center for Genomic and Computational Biology Core, Jeff Bourgeois and Dr. Kyle Gibbs, and Horner lab members for useful discussion. This work was supported by funds from Burroughs Wellcome Fund (S.M.H.), National Institutes of Health: R01AI125416, T32-CA009111 (M.J.M., M.T.S.,), and a Duke MGM SURE Fellowship (K.A.M.).

## Author contributions

Conceptualization: M.J.M., M.T.S., N.S.G., and S.M.H. Investigation: M.J.M., M.T.S., N.S.G., and M.P. Formal analysis: M.J.M., M.T.S., K.A.M., N.S.G., and S.M.H. Software: K.A.M., N.S.G., and K.Y.S. Writing – original draft: M.J.M. and S.M.H. Writing – review and editing: M.J.M., M.T.S., K.A.M., N.S.G., K.Y.S., and S.M.H. Funding acquisition: S.M.H.

## Competing interests

The authors have no competing interests to declare.

## Methods

### Cell Lines

Human hepatoma Huh7 cells and embryonic kidney 293T cells were grown in Dulbecco’s modification of Eagle’s medium (DMEM; Mediatech) supplemented with 10% fetal bovine serum (Thermo Fisher Scientific), 1X minimum essential medium non-essential amino acids (Thermo Fisher Scientific), and 25 mM HEPES (Thermo Fisher Scientific) (cDMEM). The identity of the Huh7 cells used in this study was verified by using the GenePrint STR kit (Promega) (DNA Analysis Facility, Duke University, Durham, NC, USA). Huh7 cells were a gift of Dr. Michael Gale Jr., and 293T cells (WT and PCIF1 KO) [20] were a gift of Dr. Eric Greer. All cell lines were verified as mycoplasma free by the LookOut Mycoplasma PCR detection kit (Sigma).

### IFN-β and IL-6 Treatment

All IFN-β (PBL Assay Science) and IL-6 (Sigma-Aldrich) treatments were performed at a concentration of 50 units/mL and 50 ng/mL in cDMEM, respectively.

### Plasmids

pcDNA3.1-FLAG-stm2585 (FLAG-SarA) and pcDNA3.1 empty vector plasmids [51] were gifts of Dr. Dennis Ko (Duke).

### Transfection

siRNAs directed against METTL3 (SI04317096), METTL14 (SI00459942), FTO (SI04177530), STAT3 (M-003544-02) or non-targeting AllStars negative control siRNA (1027280) were purchased from Qiagen or Dharmacon. All siRNA transfections were performed using the Lipofectamine RNAiMax reagent (Invitrogen), according to manufacturer’s instructions. siMETTL3/14 co-transfections were performed at a ratio of 1:2 siMETTL3:siMETTL14. Cells were transfected with 25 pmol of siRNA at a final concentration of 0.0125 μM. Media was changed 4 hours post-transfection, and cells were incubated for 36 hours post-transfection prior to each experimental treatment. Plasmid transfections were performed with 1 μg of DNA per single well of a 6-well plate using PEI MAX (Polysciences, INC.) according to the manufacturer’s instructions. Media was changed 4 hours post-transfection, and cells were incubated for 36 hours post-transfection prior to each experimental treatment.

### Immunoblotting

Cells were lysed in a modified radioimmunoprecipitation assay (RIPA) buffer (10 mM Tris [pH 7.5], 150 mM NaCl, 0.5% sodium deoxycholate, and 1% Triton X-100) supplemented with protease inhibitor cocktail (Sigma) and phosphatase inhibitor cocktail II (Millipore), and post-nuclear lysates were harvested by centrifugation. Quantified protein (between 5 and 15 μg) was added to a 4X SDS protein sample buffer (40% glycerol, 240 mM Tris-HCl [pH 6.8], 8% SDS, 0.04% bromophenol blue, 5% beta-mercaptoethanol), resolved by SDS/PAGE, and transferred to nitrocellulose membranes in a 25 mM Tris-192 mM glycine-0.01% SDS buffer. Membranes were stained with Revert 700 total protein stain (LI-COR Biosciences), then blocked in 3% bovine serum albumin. Membranes were incubated with primary antibodies for 2 hours at room temperature or overnight at 4°C. After washing with PBS-T buffer (1× PBS, 0.05% Tween 20), membranes were incubated with species-specific horseradish peroxidase-conjugated antibodies (Jackson ImmunoResearch, 1:5000) for 1 hour at room temperature, followed by treatment of the membrane with Clarity enhanced chemiluminescence (Bio-Rad) and imaging on an Odyssey Fc imaging system (LI-COR Biosciences). The following antibodies were used for immunoblotting: mouse anti-IFITM1 (Proteintech 60074-1-Ig, 1:1000; recognizes IFITM1 but not IFITM2 or IFITM3 [52, 53]), rabbit anti-MX1 (Abcam ab207414, 1:1000), mouse anti-ISG15 (Santa Cruz sc-166755, 1:5000), rabbit anti-EIF2AK2 (Abcam ab32506, 1:1000), rabbit anti-GBP1 (Abcam EPR8285), mouse anti-IFIT3 (Abcam ab76818), rabbit anti-METTL14 (Sigma HPA038002, 1:2500), mouse anti-METTL3 (Abnova H00056339-B01P, 1:1000), rabbit anti-FTO (Abcam EPR6895, 1:1000), mouse anti-FTO (Abcam ab92821, 1:2000), mouse anti-STAT3 (Cell Signaling Technologies 9139S, 1:2000), rabbit anti-phospho-STAT3 (Cell Signaling Technologies, 1:2000), anti-PCIF (Bethyl Laboratory, 1:1000), mouse anti-FLAG-HRP (Sigma A8592, 1:5000).

### Quantification of Immunoblots

Following imaging using the LI-COR Odyssey Fc, immunoblots were quantified using ImageStudio Lite software, and raw values were normalized to total protein (Revert 700 total protein stain) for each condition.

### RT-qPCR

Total cellular RNA was extracted using the Qiagen RNeasy kit (Life Technologies) or TRIzol extraction (Thermo Fisher Scientific). RNA was then reverse transcribed using the iScript cDNA synthesis kit (Bio-Rad) as per the manufacturer’s instructions. The resulting cDNA was diluted 1:5 in nuclease-free H2O. RT-qPCR was performed in triplicate using the Power SYBR Green PCR master mix (Thermo Fisher Scientific) and the Applied Biosystems Step One Plus or QuantStudio 6 Flex RT-PCR systems. Primer sequences for RT-qPCR are listed in Table 1.

**Table 1:**
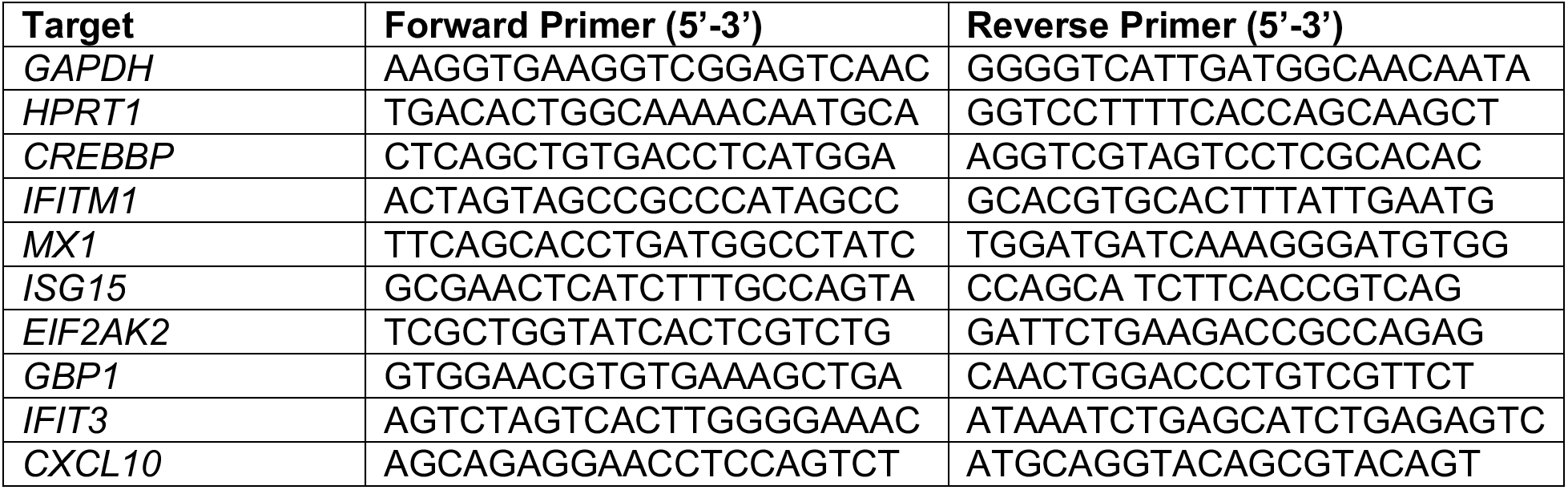
RT-qPCR Primer Sequences.

### Metabolic labeling of nascent transcripts with 4-sU

siRNA-treated cells were pulsed with 4-sU at a final concentration of 200 μM (−/+ IFN-β) for the indicated amount of time before harvest in TRIzol and RNA extraction. Purification of newly transcribed, 4-sU labeled RNA was performed as previously described [54]. Briefly, 50 μg of RNA were biotinylated in biotinylation buffer (10 mM Tris-HCl pH 7.4, 1 mM EDTA, 20 ng/μl MTSEA Biotin-XX (Biotium)) for 30 minutes at room temperature, shaking at 800 rpm. A phenol/chloroform extraction was performed, and RNA was precipitated in isopropanol for 1 hour at −20°C. Following centrifugation for 20 minutes at 12000 X g at 4°C, RNA was resuspended, and 20% of each sample was taken for input samples. The remaining RNA was then incubated with Streptavidin MyOne C1 Dynabeads (Thermo Fisher Scientific), which were prewashed with 0.1M NaCl, in streptavidin binding buffer (final concentration 5 mM Tris-HCl pH 7.4, 0.5 mM EDTA, 1M NaCl) for 30 minutes at room temperature, shaking at 800 rpm. The supernatant containing unbound “pre-existing” RNA was then collected, and the beads were washed 4X with wash buffer (100 mM Tris-HCl pH 7.4, 10 mM EDTA, 1M NaCl, 0.1% Tween-20), and each wash was collected. Two additional washes were performed, and supernatant was discarded. To elute biotinylated RNA, 100 mM DTT (freshly prepared) was added, and the supernatant was collected. This was repeated 3 times to collect all labeled RNA. Finally, both the “pre-existing” and “newly transcribed” fractions were precipitated in isopropanol for 1 hour at −20°C. Following centrifugation for 20 minutes at 12000 X g at 4°C, RNA was resuspended, and cDNA was synthesized for each RNA fraction using the iScript cDNA synthesis kit (BioRad). RT-qPCR was then performed for total and newly transcribed fractions and used to determine the relative transcription rate of genes of interest, compared to GAPDH.

### RNA-seq

Following siRNA treatment (36 hours), Huh7 cells seeded in 10-cm^2^ plates were stimulated with IFN-β or mock treated (8 hours), then harvested and RNA extraction was performed using TRIzol reagent (Thermo Fisher Scientific). Samples were then treated with Turbo DNase I (Thermo Fisher Scientific) according to manufacturer protocol and incubated at 37°C for 30 minutes, followed by phenol/chloroform extraction and ethanol precipitation overnight. RNA concentrations were then normalized. Sequencing libraries were prepared using the KAPA Stranded mRNA-Seq Kit (Roche) and sequenced on an Illumina HiSeq 4000 with 100 bp paired-end reads by the Duke University Center for Genomic and Computational Biology.

Reads were aligned using Salmon [55] to the human reference transcriptome using default parameters (hg19). Differential gene expression between infected and uninfected samples was compared using DESeq2 [56]. Gene ontology analyses were generated using the PANTHER Classification System’s statistical enrichment test with gene symbols and fold change values [57]. The heatmaps were generated using Python scripts. We compared the effects of IFN-B and mock treatment in cells transfected with siCTRL to determine the most highly induced interferon stimulated genes based on adjusted P-value < 0.01 and a basemean > 50. To determine the effect of IFN-β treatment with FTO depletion, the log-transformed adjusted P-value and fold changes of the previously determined highly induced genes were plotted from the IFN-β and mock treatments using Python scripts. Heatmaps of the 50 most induced ISGs and pro-inflammatory chemokines and cytokines were generated using Python scripts.

### Data Availability

All raw data from RNA-seq are available through GEO (accession number: GSE180663).

### Code Availability

All RNA-seq analysis scripts are open-source or online on Github (https://github.com/kristen-a-murphy/McFadden_siFTO_RNA-seq).

## Supplemental Information

Table S1: RNA-seq analysis of gene expression changes following IFN-β treatment and FTO depletion.

- Table S1.1: siCTRL IFN / siCTRL Mock
- Table S1.2: siFTO IFN / siCTRL IFN
- Table S1.3: siFTO Mock / siCTRL Mock

Table S2: Gene Ontology analysis of RNA-seq data.

- Table S2.1: Enriched GO categories for differentially expressed genes (siFTO Mock / siCTRL Mock)
- Table S2.2: Enriched GO categories for differentially expressed genes (siFTO IFN / siCTRL IFN)

## Notes

### Competing Interest Statement

The authors have declared no competing interest.

